# The TAF10-containing TFIID and SAGA transcriptional complexes are dispensable for early somitogenesis in the mouse embryo

**DOI:** 10.1101/071324

**Authors:** Paul Bardot, Stéphane D. Vincent, Marjorie Fournier, Alexis Hubaud, Mathilde Joint, László Tora, Olivier Pourquié

**Affiliations:** Institut de Génétique et de Biologie Moléculaire et Cellulaire, Illkirch, France; Centre National de la Recherche Scientifique, UMR7104, Illkirch, France; Institut National de la Santé et de la Recherche Médicale, U964, Illkirch, France; Université de Strasbourg, Illkirch, France

**Keywords:** RNA polymerase II, TATA binding protein, presomitic mesoderm, paraxial mesoderm, conditional knock out, proteomic

## Abstract

During development, tightly regulated gene expression programs control cell fate and patterning. A key regulatory step in eukaryotic transcription is the assembly of the pre-initiation complex (PIC) at promoters. The PIC assembly has mainly been studied *in vitro*, and little is known about its composition during development. *In vitro* data suggests that TFIID is the general transcription factor that nucleates PIC formation at promoters. Here we show that TAF10, a subunit of TFIID and of the transcriptional co-activator SAGA, is required for the assembly of these complexes in the mouse embryo. We performed *Taf10* conditional deletions during mesoderm development and show that *Taf10* loss in the presomitic mesoderm (PSM) does not prevent cyclic gene transcription or PSM segmental patterning, while lateral plate differentiation is profoundly altered. During this period, global mRNA levels are unchanged in the PSM, with only a minor subset of genes dysregulated. Together, our data demonstrate that the TAF10-containing canonical TFIID and SAGA complexes, are dispensable for early paraxial mesoderm development, arguing against the generic role in transcription proposed for these fully assembled holo complexes.

## Introduction

In mouse, the posterior part of the paraxial mesoderm, called presomitic mesoderm (PSM), generates a pair of somites every two hours and plays crucial roles during vertebrate elongation (Pourquié, 2011). This rhythmic process is under the control of a clock characterized by periodic waves of transcription of cyclic genes sweeping from the posterior to the anterior PSM (Hubaud and Pourquié, 2014). In the anterior PSM, the clock signal is converted into a stripe of expression of specific segmentation genes that defines the future somite. This periodic transcription initiation associated to the segmentation clock oscillations in the PSM offers a unique paradigm to study transcriptional regulation in development.

During embryogenesis, gene expression is regulated by a combination of extracellular signals triggering intracellular pathways, which converge towards the binding of transcription factors to enhancers and promoters. These interactions lead to the assembly of the transcriptional machinery. In non-plant eukaryotes, there are three RNA polymerases able to transcribe the genome and among them, RNA polymerase II (Pol-II) is responsible for the production of mRNA and some of the non-coding RNAs (Levine et al., 2014).

Transcription initiation requires the assembly of the Pre-Initiation Complex (PIC) that allows the correct positioning of Pol-II on the promoter and the consequent RNA synthesis (Sainsbury et al., 2015). TFIID is the first element of the PIC recruited to active promoters. In its canonical form in higher eukaryotes, it is composed of TATA binding protein (TBP) and 13 TBP-Associated Factors (TAFs) and is involved in the correct positioning of Pol-II on the transcription start site. While TBP is also part of Pol-I and Pol-III transcription complexes, the TFIID-TAFs are specific for Pol-II transcription machinery. Among the metazoan TAFs, TAF9, TAF10 and TAF12 are also shared by Spt-Ada-Gcn5-acetyl transferase (SAGA) complex, a transcriptional co-activator conserved from yeast to human (Spedale et al., 2012). SAGA exhibits HAT activity at the promoters and also deubiquitylates histone H2Bub in gene bodies (Bonnet et al., 2014; Wang and Dent, 2014; Weake et al., 2011).

Several structural TAFs, including TAF10, share a Histone Fold Domain (HFD) involved in their dimerization with specific partners: TAF10 heterodimerizes with TAF3 or TAF8 within TFIID and with SUPT7L/ST65G within SAGA (Leurent et al., 2002; Soutoglou et al., 2005). Nuclear import of TAF10 is absolutely dependent on heterodimerization with its partners since TAF10 does not have a nuclear localization signal (NLS) (Soutoglou et al., 2005).

TAF10 does not exhibit any enzymatic activity but serves as an interface allowing interaction with other TAFs (Bieniossek et al., 2013; Trowitzsch et al., 2015) or transcription factors such as the human estrogen receptor α (Jacq et al., 1994) or mouse GATA1 (Papadopoulos et al., 2015). In HeLa cells, only 50% of the TFIID complexes contain TAF10 (Jacq et al., 1994). A lower proportion of TFIID complexes lacking TAF10 has also been observed in F9 cells (Mohan et al., 2003), but their functionality is unknown. The structure of TFIID in the absence of TAF10 is unclear. Only partial TFIID sub-complexes, not associated with TBP, were detected in undifferentiated and retinoic acid (RA) differentiated *Taf10* mutant F9 cells (Mohan et al., 2003), while lack of TFIID was observed in *Taf10* mutant liver cells (Tatarakis et al., 2008). SAGA was not investigated in these experiments (Mohan et al., 2003; Tatarakis et al., 2008). Altogether, these data support the idea that TFIID composition can vary, as also suggested by the existence of TAF paralogs and/or tissue specific TAFs (Goodrich and Tjian, 2010; Müller et al., 2010).

The diversity in TFIID’s composition may have functional consequences. Whereas TAF10 is crucial for survival and proliferation of F9 cells, it is dispensable for their differentiation into primitive endoderm (Metzger et al., 1999). *Taf10* mutation in mouse leads to embryonic lethality shortly after implantation (Mohan et al., 2003). Interestingly, while inner cell mass cells die by apoptosis, trophoectodermal cells survive, although Pol-II transcription is greatly reduced (Mohan et al., 2003). *Taf10 c*onditional deletion in skin or liver has shown that TAF10 is required for transcription in the embryo, but not in the adult (Indra et al., 2005; Tatarakis et al., 2008). Altogether, these data indicate that TAF10 requirement depends on the cellular and developmental context.

In this study, we aimed to closely analyse TAF10 requirement and its role in transcription during mouse development, and to examine the composition of TFIID and SAGA in the absence of TAF10 in embryonic tissues *in vivo*. We performed immunoprecipitations coupled to mass spectrometry analyses on embryonic lysates. We show that, in the mouse embryo, absence of TAF10 severely impairs TFIID and SAGA assembly. In order to get insights into the functional importance of TAF10 during development, we focused on paraxial mesoderm dynamic differentiation by carrying out a *Taf10* conditional deletion in the mesoderm using the *T-Cre* line (Perantoni, 2005). While loss of *Taf10* eventually led to growth arrest and cell death around E10.5, we identified a time window during which the dynamic transcription of cyclic genes is still maintained in the absence of detectable TAF10 protein. Microarray analysis of mutant PSM revealed that Pol-II transcription is not globally affected in this context, although expression of some genes, such as genes encoding cell cycle inhibitors, is up regulated.

## Results

### TAF10 is ubiquitously expressed in the nucleus of embryonic cells at E9.5

*Taf10* is ubiquitously expressed in the mouse embryo at E3.5, E5.5 and E7.5 but with more heterogeneity at E12.5 (Mohan et al., 2003). *W*hole-mount *in situ* hybridization (WISH) analyses suggest that *Taf10* is also ubiquitously expressed at E8.5 and E9.5 (Fig. S1A,B). TAF10 protein is ubiquitously expressed in the posterior part of the embryo (Fig. S1C, Fig. S2) and no heterogeneity was observed between E9.5 and E10.5. Competition with the peptide used to raise the anti-TAF10 antibody (Mohan et al., 2003) confirms that TAF10 localization is specific since the TAF10 signal, but not the Myogenin signal, is lost under these conditions (Fig. S1D,H). Altogether these results indicate that TAF10 protein is ubiquitously expressed in cell nuclei between E8.5 and E10.5.

### Induced ubiquitous deletion of *Taf10* leads to growth arrest at E10, but does not impair transcription at E9.5

In order to analyse the effects of TAF10 absence on development, we performed a *Taf10* inducible ubiquitous deletion using the *R26*^CreERT2^ line (Ventura et al., 2007). This strategy deletes the exon 2 of *Taf10* that encodes for part of the HFD (Mohan et al., 2003). Since the exon 3 is out of frame, this deletion is expected to produce a 92 amino-acids-truncated protein without HFD (Fig. S3D). Since the HFD is required for heterodimerization and integration of TAF10 into TFIID and SAGA (Leurent et al., 2002; Soutoglou et al., 2005), this potential truncated protein cannot be integrated into mature SAGA or TFIID complexes. Tamoxifen was injected intraperitoneally at E7.5 and Cre recombination was followed by the activity of the Cre reporter allele *R26*^R^ (Soriano, 1999). Complete Cre recombination is observed at E9.5 (Fig. 1A,B). The development of *R26*^CreERT2/+^;*Taf10*^flox/flox^ (*R26Cre;Taf10*) mutant embryos was arrested at E9.5 as embryos do not further develop when recovered at E10.5 and E11.5 (Fig. 1D,F). The normal development of *R26*^R/+^;*Taf10*^flox/flox^ embryos littermates (Fig. 1C,D) confirmed that tamoxifen injection at E7.5 does not induce secondary defects.

**Fig. 1.**
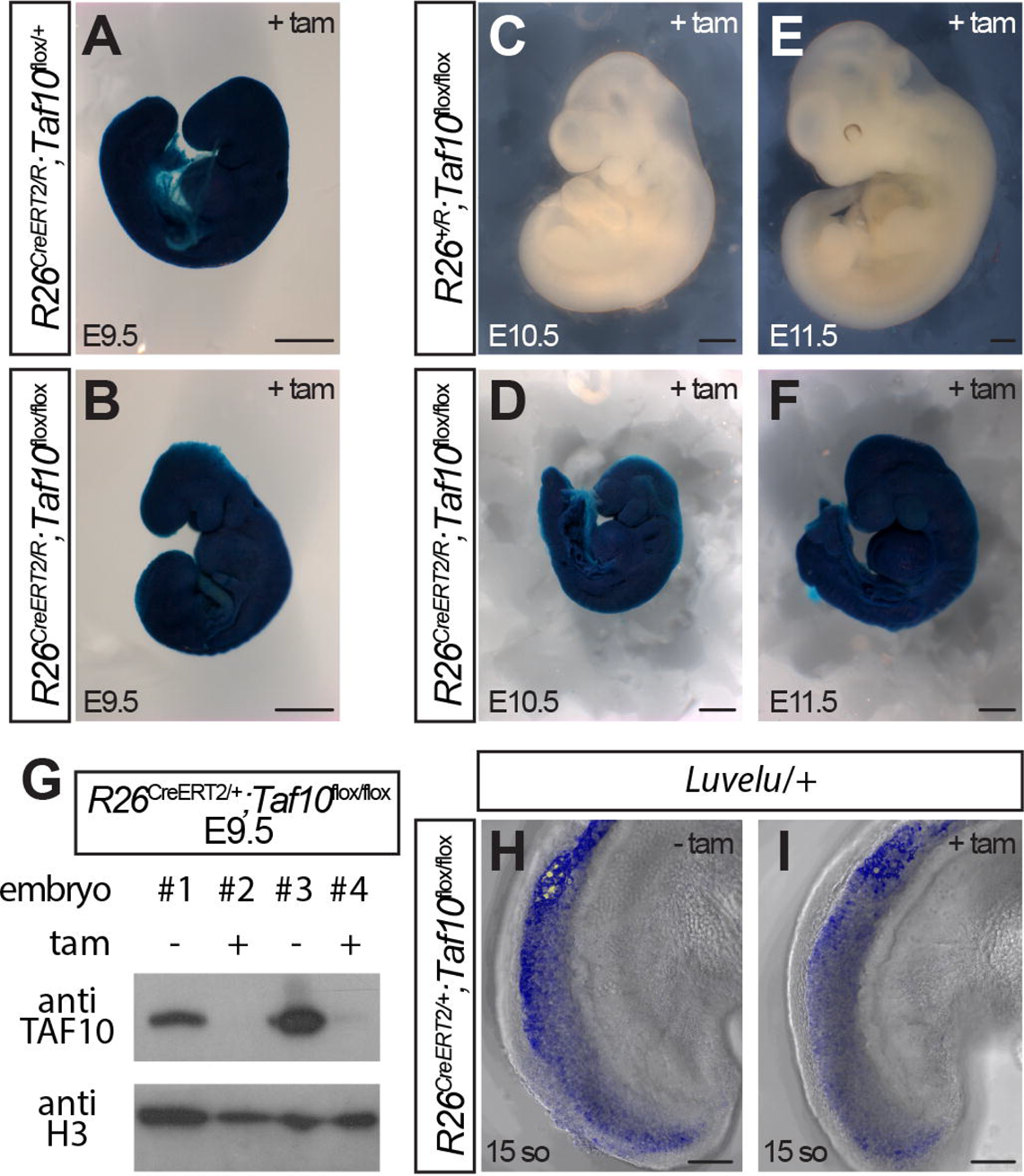
Efficient ubiquitous deletion of *Taf10* in E9.5 *R26Cre*;*Taf10* mutant embryos. (A-F) Whole-mount Xgal staining of E9.5 *R26*^CreERT2/R^;*Taf10*^flox/+^ (A) or E10.5 (C) and E11.5 (E) *R26*^+/R^;*Taf10*^flox/flox^ control embryos and E9.5 (B), E10.5 (D) and E11.5 (F) *R26*^CreERT2/R^;*Taf10*^flox/flox^ mutant embryos after tamoxifen treatment at E7.5 (+tam). (G) Western blot analysis of E9.5 *R26Cre*;*Taf10* whole embryos treated (+) or not (-) with tamoxifen at E7.5 with anti-TAF10 (top) and anti-H3 (bottom) antibodies. (H-I) Confocal z-stack images projection of E9.25 *R26*^CreERT2/+^;*Taf10*^flox/flox^;*Luvelu/+* non treated (H) or treated (I) embryos with tamoxifen (+tam). Scale bars in A-F and H-I represent 500 µm, and 100 µm, respectively tam; tamoxifen.

Whereas no TAF10 protein was detected by western blot at E9.5 after tamoxifen injection at E7.5 (Fig. 1G), TAF10 was still present, albeit at lower levels, at E8.5 (Fig. S3E). This observation is in agreement with a previous study in which TAF10 protein was still detected one day after induction of its depletion (Metzger et al, 1999). Since our goal is to assess TFIID and SAGA composition in absence of TAF10, we performed our analyses at E9.5.

In order to assess transcription initiation *in vivo*, we used the *Luvelu* line (Aulehla et al., 2008) that allows visualization of the dynamic waves of *Lfng* transcription occurring every 2 hours in the posterior PSM. This line contains the promoter and 3’-UTR destabilizing sequences of the cyclic gene *Lfng* (Cole et al., 2002; Morales et al., 2002), coupled to the coding sequences of a Venus-PEST fusion. *Luvelu* expression is not affected in the absence of TAF10 at E9.5 (Fig. 1H,I), clearly indicating that transcription initiation still occurs in the *R26Cre;Taf10* mutant embryos, at least in the PSM. Altogether, these results show that in mutants in which *Taf10* deletion is induced at E7.5, no TAF10 protein is detected in the PSM at E9.5 yet periodic gene transcription in the PSM is not affected.

### Analyses of TFIID and SAGA composition in the absence of TAF10 in the mouse embryo

Next, we set out to analyse TFIID and SAGA composition by mass spectrometry in E9.5 mouse embryos, when no TAF10 protein is detected. To purify these complexes, we collected E9.5 embryos from *R26*^CreERT2/CreERT2^;*Taf10*^flox/flox^ x *Taf10*^flox/flox^ crosses, treated (mutant) or not (control) with tamoxifen at E7.5. Complete *Taf10* deletion was assessed by PCR (data not shown) and western blot analysis that confirmed the absence of detectable full-length TAF10 protein (Fig. 2A). Interestingly, in whole cell extracts from mutants, expression of TBP, TAF4A, TAF5 and TAF6 is not affected, whereas expression of TAF8, the main TFIID partner of TAF10, is strongly decreased (Fig. 2A), indicating that the TAF8-TAF10 interaction is required for the stabilization of TAF8. We then compared TFIID and SAGA composition in the presence or absence of TAF10 by performing immunoprecipitations (IPs) of different TFIID and SAGA subunits using anti-TBP, or anti-TAF7 antibodies (for TFIID) and with anti-TRRAP or anti-SUPT3 (for SAGA). Composition of the immuno-precipitated complexes was analysed by mass spectrometry (Table S1). The normalized spectral abundance factor (NSAF) values were calculated to compare between control and *Taf10* mutant samples (Zybailov et al., 2006).

**Fig. 2.**
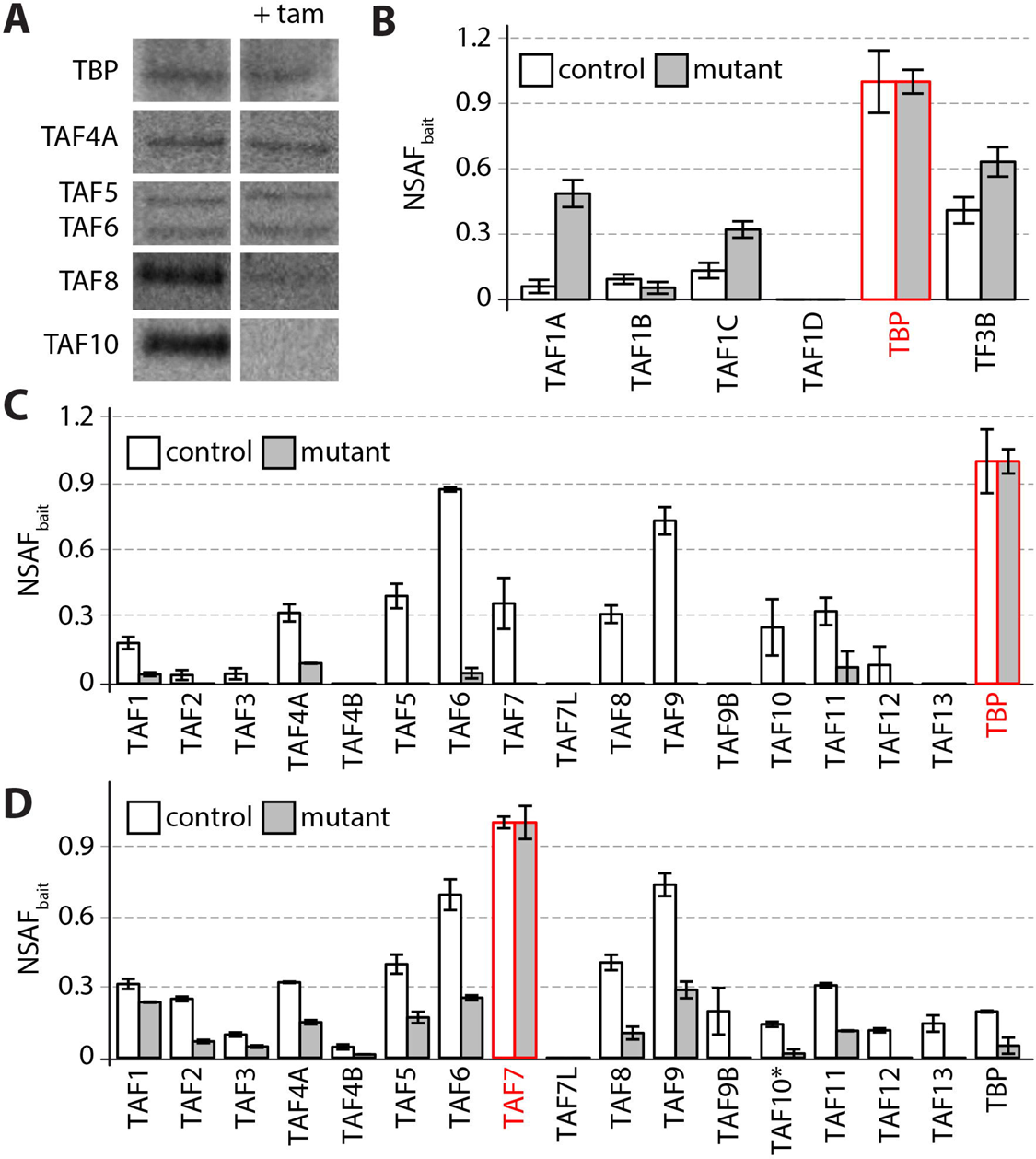
TFIID assembly defect in *R26Cre;Taf10* mutant embryos. (A) Western blot analysis of the expression of TBP, TAF4A, TAF5, TAF6, TAF8 and TAF10 from whole cell extracts of E9.5 *R26Cre;Taf10* control (left, untreated) or mutant (right, treated with tamoxifen at E7.5) embryos. (B) TBP NSAFbait values for SL1 complex subunits (TAF1A, TAF1B, TAF1C, TAF1D and TBP) and TF3B-TBP complex. (C-D) NSAFbait values for TFIID subunits of TBP-IP (C) and TAF7-IP (D). Bait proteins are indicated in red. Control and mutant IPs are indicated in white and grey, respectively. TAF10* corresponds to the full-length TAF10 protein. tam; tamoxifen

In control embryos, the full-length TAF10 protein is represented by 4 peptides (Fig. S4A). In mutant embryo samples, no TAF10 peptides were detected in TBP- and TRRAP-IPs. In contrast, in TAF7- and SUPT3-IPs, we detected significant amounts (albeit reduced compared to control) of the TAF10 N-terminal peptide (peptide #1) (Fig. S4B,C). The *Taf10*^flox^ conditional mutation deletes the exon 2 resulting in an out of frame fusion of exon 1 to exon 3 leading to premature truncation of the protein TAF10. This deletion is thus expected to produce a truncated N-terminal fragment of TAF10 containing peptide #1, but not the other peptides (Fig. S4D). The fact that no TAF10 peptides are detected in TBP- and TRRAP-IPs suggests that the truncated N-terminal peptide remaining in the mutant cannot participate to fully assembled TFIID or SAGA complexes. In addition, importantly, no TFIID subunits could be immunoprecipitated from murine *R26*^CreERT2/R^;*Taf10*^flox/flox^ ES cells, after 4-hydroxy tamoxifen treatment, with an antibody that recognizes the N terminal part of the TAF10 protein (Fig. S3B) and is able to immunoprecipitate the TFIID complex (Fig. S5A,B), showing that the truncated peptide is not part of a fully assembled TFIID complex. No conclusion could be drawn for the SAGA complex since this anti Nterm-TAF10 antibody is not able to co-immunoprecipitate any of the mouse SAGA subunits even in control conditions (Fig. S5C). These data are consistent with the fact that the mutant truncated protein does not contain the HFD (Soutoglou et al., 2005). Thus, for further analyses and to score only the full-length protein, we took into account peptides #2 to #4 that should be absent from the full-length TAF10 protein after deletion of the genomic sequences (TAF10*, Fig. 2D and Fig. 3C, and Fig. S4A,D) for TAF7-IPs (Fig. 2D) and SUPT3-IPs (Fig. 3C). Data for TAF7- and SUPT3-IPs taking into account all the TAF10 peptides are shown in Fig. S6.

TBP is also part of SL1 and TFIIIB complexes, involved in Pol-I and Pol-III transcription, respectively (Vannini and Cramer, 2012). Importantly, TAF10 absence does not perturb the interaction of TBP with its non-TFIID partners, highlighting the lack of non-specific effect (Fig. 2B). In *Taf10* mutant embryos, we observed an increased interaction between TBP and the larger SL1 subunits; TAF1A and TAF1C, suggesting that TBP may be redistributed in Pol-I TAF-containing complexes in TAF10 absence. This is consistent with the observation that there is no free TBP in the cells (Timmers and Sharp, 1991). In control TBP- and TAF7-IPs, all the canonical TFIID subunits were detected (Fig. 2C,D). Interestingly, in *Taf10* mutant embryos, TBP-IP reveals that TBP is mostly disengaged from TFIID as only a few TAFs co-immunoprecipitate with TBP in low amounts (Fig. 2C). This TFIID dissociation is also observed in the TAF7-IP in absence of TAF10 (Fig. 2D). Surprisingly however, due to the very efficient TAF7-IP (Table S1), we can still detect residual canonical TFIID complexes (Fig. 2D). It is important to note that, if the anti-TAF7 antibody is able to co-immunoprecipitate most of the TAFs, TAF9B, TAF12 and TAF13 are not detected in the mutant, further supporting the conclusion that TAF10 absence strongly affects TFIID assembly.

In order to assess SAGA composition, we performed IPs against two SAGA subunits: SUPT3 and TRRAP. TRRAP is also a member of the chromatin remodelling complex TIP60/NuA4 (Sapountzi and Côté, 2011). As the interactions between TRRAP and TIP60/NuA4 subunits were not affected (Fig. 3A), we conclude that TAF10 absence does not interfere with the interactions between TRRAP and its non-SAGA partners. In both mutant TRRAP-IP (Fig. 3B) and SUPT3-IP (Fig. 3C), we observed a dramatic reduction in the amount of SAGA subunits co-immunoprecipitated, clearly showing a defect in the assembly of SAGA. Contrary to TAF7-IP, we were not able to detect any residual canonical SAGA complexes in the mutant samples.

**Fig. 3.**
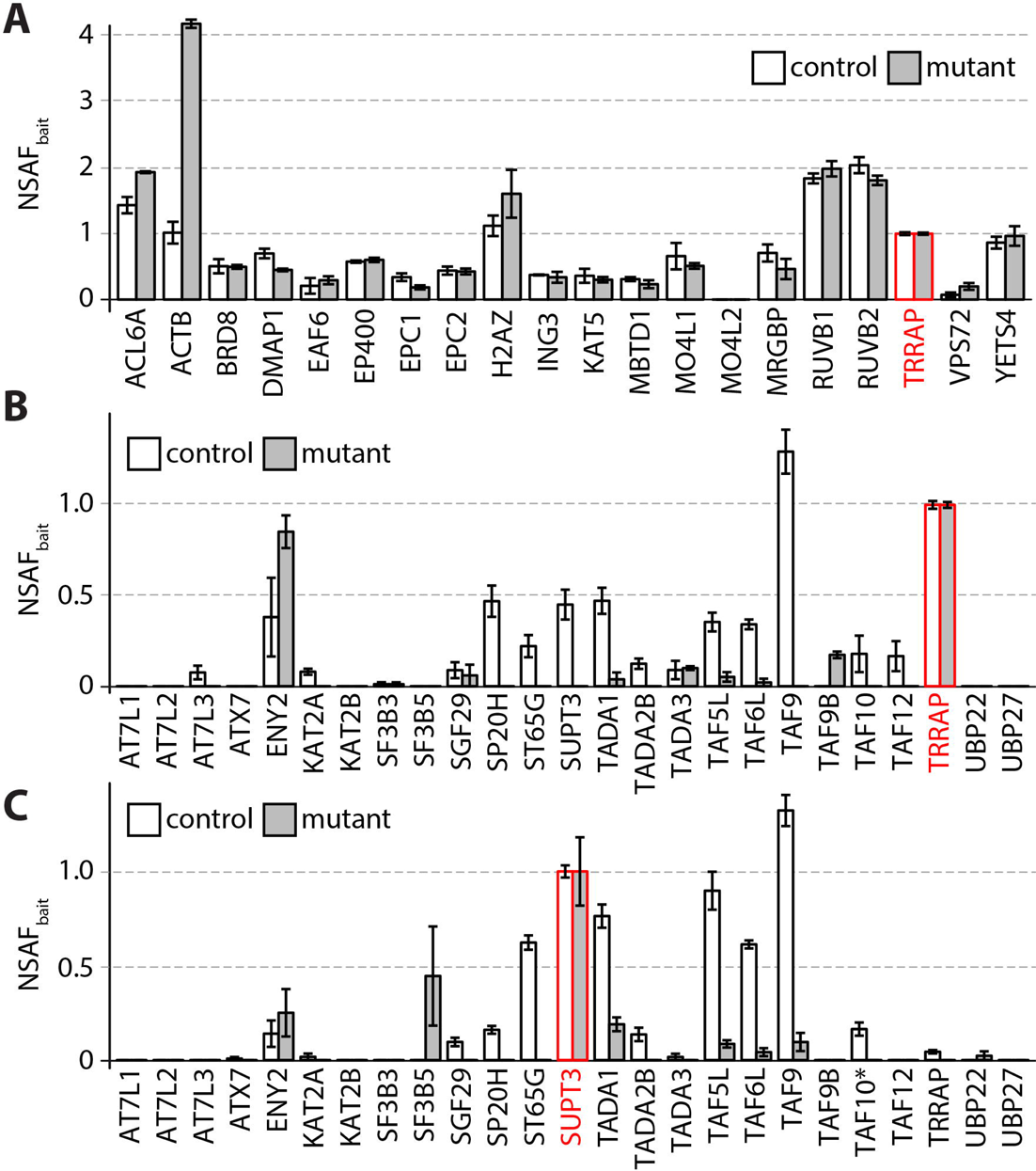
SAGA assembly defect in *R26Cre;Taf10* mutant embryos. (A) NSAFbait values for TIP60/NuA4 complex subunits of TRRAP-IP from control (white) and mutant (grey) extracts. (B-C) NSAFbait values for SAGA subunits of TRRAP-IP (C) and SUPT3-IP (D) from control (white) and mutant (grey) extracts. Bait proteins are indicated in red. TAF10* corresponds to the full-length TAF10 protein.

Altogether, these results strongly suggest that TAF10 is crucial for the assembly of both TFIID and SAGA in the mouse embryo, since the formation of both complexes is seriously impaired in *R26Cre;Taf10* mutant embryos.

### *Taf10* conditional deletion in the paraxial mesoderm

Our next goal was to analyse TAF10 requirement for transcription during development. Somitogenesis is a dynamic developmental process in vertebrate embryos relying on periodic transcriptional waves sweeping from posterior to anterior in the PSM (Hubaud and Pourquié, 2014). As described above, the dynamic expression of the *Luvelu* cyclic reporter is not affected in the PSM of E9.5 *R26Cre;Taf10* mutant embryos (Fig. 1H,I). We next carried out a *Taf10* conditional deletion of in the PSM using the *T-Cre* line (Perantoni, 2005). This line expresses Cre in the primitive streak under the control of 500-bp *T* promoter sequences (Clements et al., 1996) leading to efficient recombination in the mesoderm before E7.5, including in paraxial mesoderm progenitors (Perantoni, 2005). *Taf10* conditional deletion is embryonic lethal as no *T-Cre/+*;*Taf10*^flox/flox^ (*T-Cre;Taf10*) mutants could be recovered at birth (data not shown). At E9.25, control and *T-Cre;Taf10* mutant embryos are very similar, except that some mutant embryos show a curved trunk (Fig. 4A,B). At E10.25, *T-Cre;Taf10* mutant embryos exhibit normal anterior development but show an apparent growth arrest of the trunk region, an helicoidal trunk lacking limb buds (Fig. 4C,D) and a degeneration of the allantois and placenta (data not shown). Whereas at E9.25 mutant and control somites were morphogically similar (Fig. 4A,B), E10.25 mutant somites were much smaller compared to the controls (Fig. 4C,D). Similar observations were made using the *Hes7-Cre* line (data not shown) that has a similar recombination pattern in the mesoderm (Niwa et al., 2007). Lysotracker Red staining indicates that there is no obvious cell death in the mutants at E9.25 (Fig. 4E,F). Recombination in the mesoderm is efficient as shown by the profile of activation of the Cre reporter allele *R26*^R^ at E8.75 (Fig. 4G,H). The full-length TAF10 protein expression could not be detected in the mesoderm of mutant embryos as early as E8.5 (Fig. S7 and 4I-L), including the PSM at E9.5 (Fig. 4I,J), while it is detected in the ectoderm. TAF10 expression was mosaic in the mutant neural tube (NT) which shares common progenitors with the mesoderm (Gouti et al., 2014; Tzouanacou et al., 2009). Surprisingly, these data show that there is a time window around E9.5 when embryonic development is not affected upon TAF10 depletion, except for the absence of limb buds, prior to an apparent growth arrest and decay at E10.5.

**Fig. 4.**
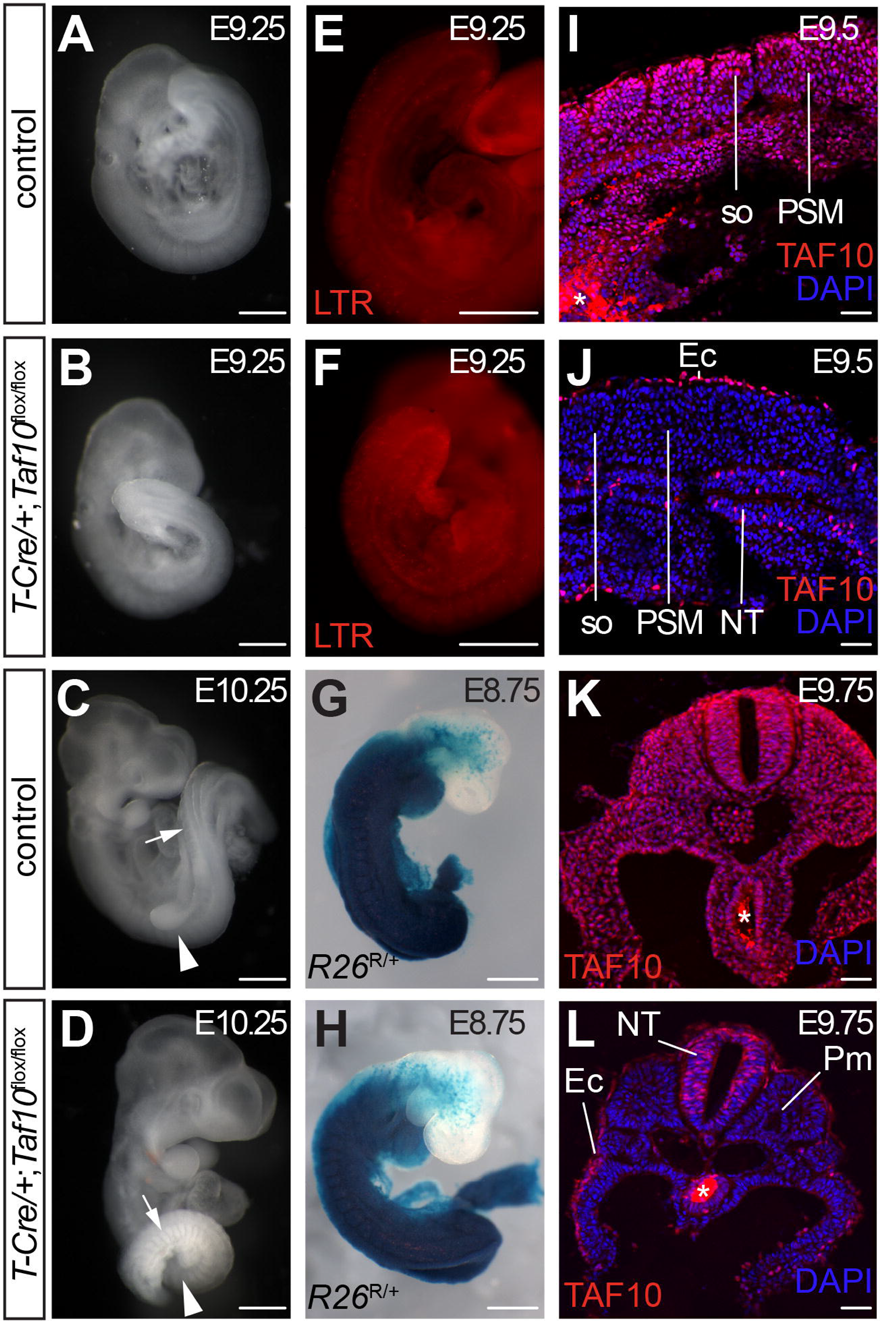
Efficient *Taf10* conditional deletion in the paraxial mesoderm. (A-C) Whole-mount right sided view of control (A,C) and *T-Cre;Taf10* mutant (B,D) E9.25 embryos (A,B) and E10.25 (C,D). The arrowhead in C,D indicates the position of the forelimb bud that is absent in the mutant, the arrow indicates the somites. (E,F) Cell death assay by LysoTracker Red staining of E9.25 control (E) and *T-Cre;Taf10* mutant (F) embryos. (G,H) Whole-mount X gal stained E8.75 *T-Cre*/+;*R26*^R/+^ control (G) and *T-Cre*/+;*R26*^R/+^;*Taf10*^flox/flox^ mutant (H) embryos showing the efficient early recombination within the paraxial mesoderm. (I-L) DAPI counterstained TAF10 immunolocalization on sagittal (I,J) and transverse (K,L) sections from E9.5 (I,J) and E9.75 (K,L) control (I,K) and *T-Cre;Taf10* mutant (J,L) embryos. so; somites, PSM; presomitic mesoderm, NT; neural tube, Ec; ectoderm, Pm; paraxial mesoderm. The asterisk (*) in (K,L) indicates background due to secondary antibody trapping in the endoderm lumen. Scale bars in A-H and I-L represent 500 µm and 50 µm, respectively.

### Absence of TAF10 in the PSM does not affect somitogenesis at E9.5

To get more insight into somitogenesis, we compared the somite numbers between the different genotypes at E9.5 (Fig. 5A). Although no significant statistical differences could be detected, mutant embryos tend to have half a somite less than the other genotypes. This could be explained by a slowing down of somitogenesis at late E9.5 stage.

**Fig. 5.**
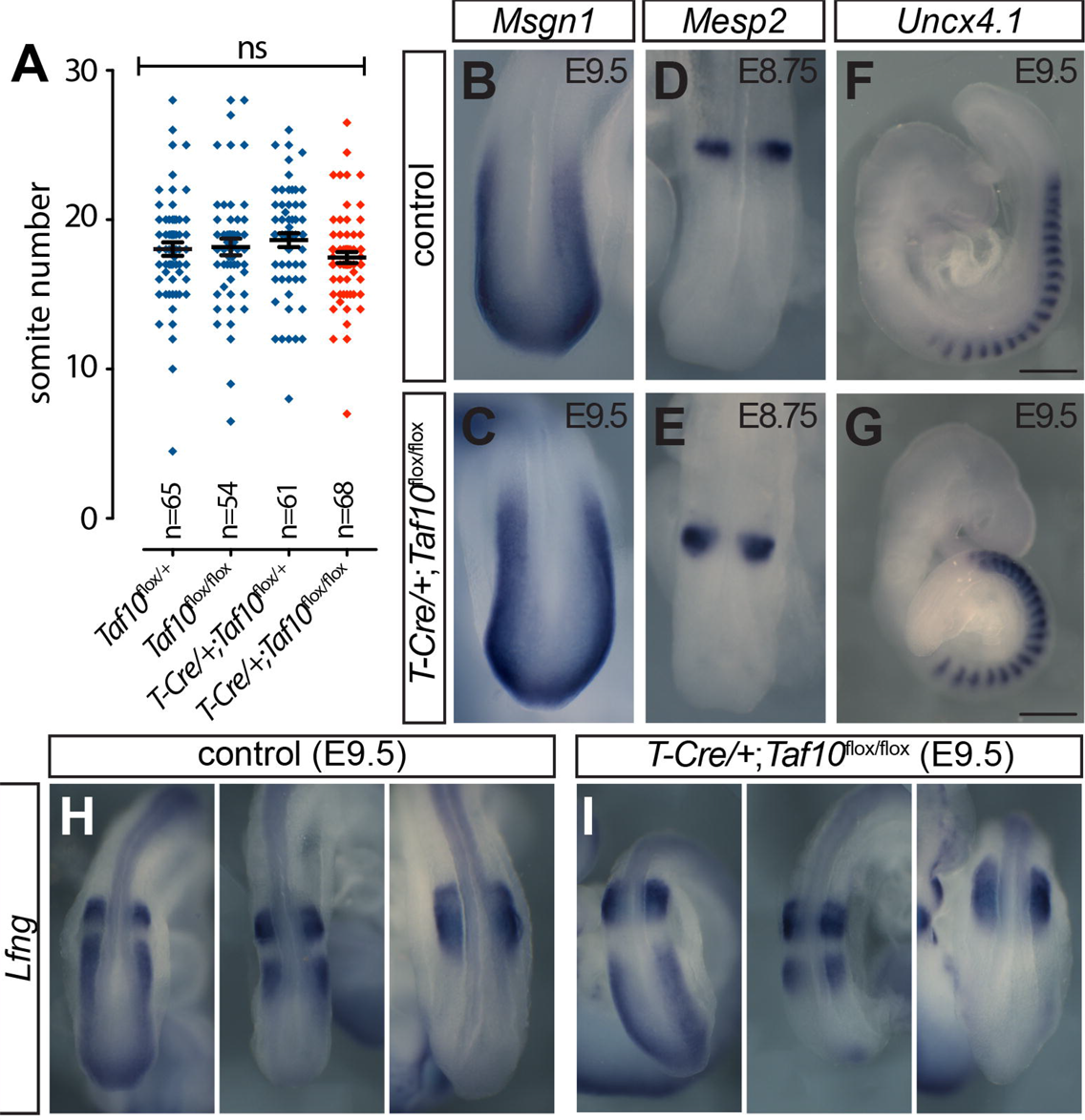
Absence of TAF10 in the PSM does not affect segmentation. (A) Somites number quantification (one-way ANOVA, ns; non significant). The error bars indicate the s.e.m. and the middle bar indicates the mean. (B-I) Whole-mount *in situ* hybridization of E9.5 (B,C,F,G,H,I) and E8.75 (D,E) control (B,D,F,H) and *T-Cre*/+;*Taf10*^flox/flox^ mutant (C,E,G,I) embryos using posterior PSM marker *Msgn1* (B,C), segmentation gene *Mesp2* (D,E), caudal somite marker *Uncx4.1* (F,G) and cyclic gene *Lfng* (H,I) probes. Dorsal tail tip (B-E,H,I) or right-lateral views (F,G) are presented. Scale bars in B-E,H-I and F-G represent 100 µm and 500 µm, respectively.

We next analysed the expression of specific PSM markers using WISH. Expression of the posterior PSM marker *Msgn1* (Wittler et al., 2007) (Fig. 5B,C), of the segmentation gene *Mesp2* (Saga et al., 1997) (Fig. 5D,E) or of the caudal somite marker *Uncx4.1* (Neidhardt et al., 1997) (Fig. 5F,G) is not affected in absence of TAF10. WISH of cyclic genes of the Notch (*Lfng* (Forsberg et al., 1998; McGrew et al., 1998) and *Hes7* (Bessho et al., 2003), Fig. 5H,I and S8A,B), Wnt (*Axin2* (Aulehla et al., 2003), Fig. S8C,D) or FGF (*Snai1* (Dale et al., 2006), Fig. S8E,F) pathways revealed that the different phases of expression could be observed in *T-Cre;Taf10* mutant embryos. Altogether, the rhythmic transcription of the cyclic genes in absence of TAF10 suggests that active transcription proceeds normally in the PSM of mutant embryos.

### Absence of TAF10 differentially affects the mesoderm derivatives

Limb bud outgrowth requires signals such as FGF8 from the Apical Ectodermal Ridge (AER) which controls proliferation of the underlying mesenchyme derived from the lateral plate mesoderm (LPM) (Zeller et al., 2009). On E10.25 transverse sections from control embryos, mesodermal nuclei (including in the LPM) are regularly shaped (Fig. 6A,C,E). However, in *T-Cre;Taf10* mutants (Fig. 6B), the paraxial mesoderm nuclei appear normal (Fig. 6D) while in the LPM (and in the intermediate mesoderm (data not shown)), we observed massive nuclear fragmentation characterized by the presence of pyknotic nuclei (Fig. 6F). Since we did not observe any difference in the efficiency in TAF10 protein depletion between the paraxial mesoderm and the LPM as early as E8.5 (Fig. S7), these data indicate that the LPM is more sensitive to *Taf10* loss than the paraxial mesoderm.

**Fig. 6.**
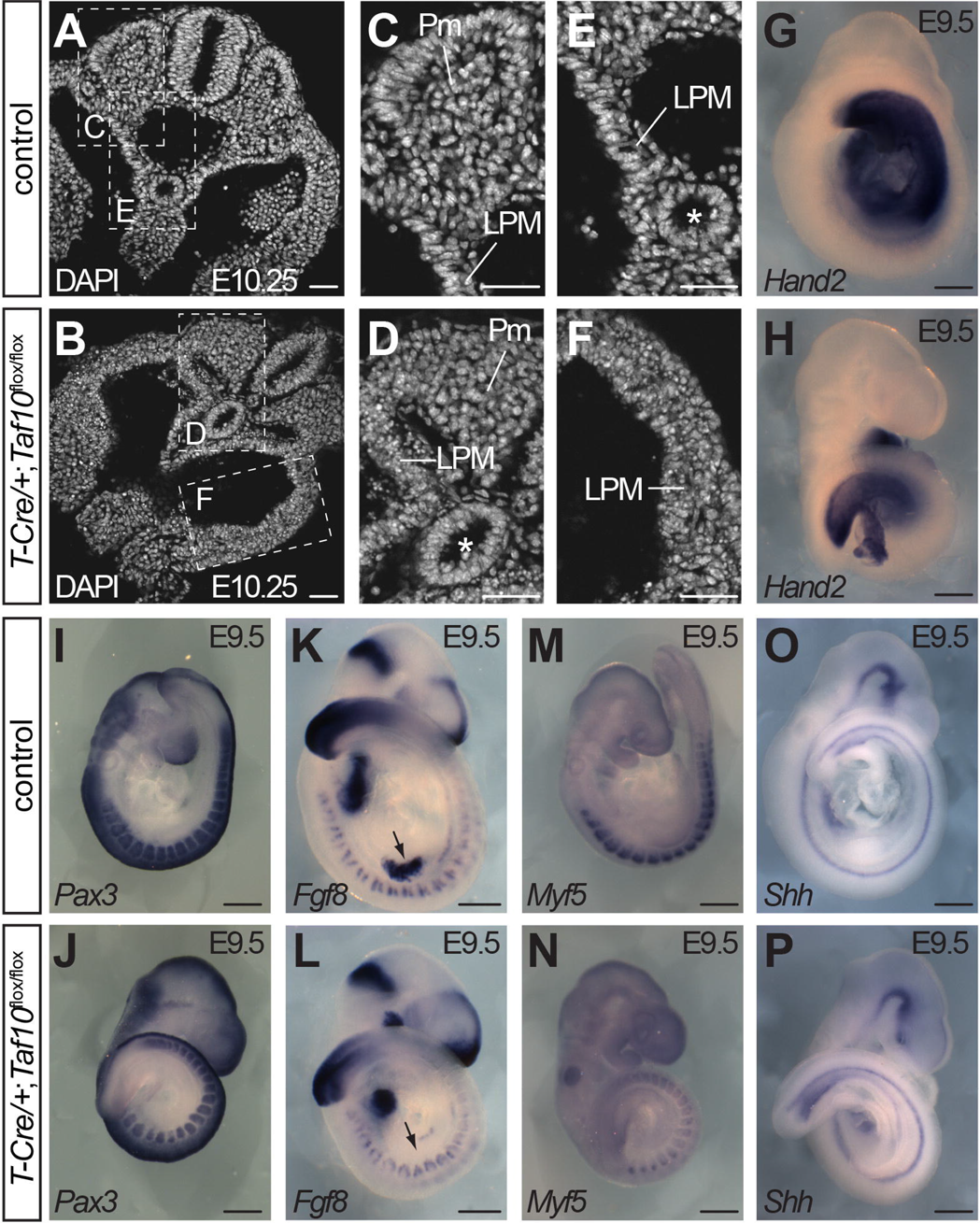
Absence of TAF10 differentially affects the different types of mesoderm. (A-F) DAPI-stained transversal sections of E10.25 control (A, magnifications in C,E) and *T-Cre*/+;*Taf10*^flox/flox^ mutant (B, magnifications in D,F) embryos showing nuclear fragmentation in LPM but normal nuclear morphology in the paraxial mesoderm. (G-P) Whole-mount *in situ* hybridization of E9.5 control (G,I,K,M,O) and *T-Cre*/+;*Taf10*^flox/flox^ mutant (H,J,L,N,P) embryos using *Hand2* (G,H), *Pax3* (I,J), *Myf5* (K,L), *Fgf8* (M,N) and *Shh* (O,P). The arrow in M,N indicates the Apical Ectodermal Ridge. Pm; paraxial mesoderm, LPM; lateral plate mesoderm, the asterisks (*) in D and E indicate the endoderm. Scale bars in A-F and G-P represent 50 µm and 500 µm, respectively.

We carried out WISH in order to test whether *Taf10* loss differentially affects the expression of specific markers of the different types of mesoderm. Expression of the LPM marker *Hand2* (Fernandez-Teran et al., 2000) is clearly down-regulated in the mutants (Fig. 6G,H). Similar observations were made with *Prdm1* that is expressed in the growing mesenchyme during limb bud outgrowth (Vincent et al., 2005) (data not shown). The absence of *Fgf8* induction in the presumptive AER in E9.5 *T-Cre;Taf10* mutant embryos (Fig. 6K,L) indicates that the LPM defect is early and probably precedes the cell death in this tissue since no obvious cell death could be detected at E9.25 (Fig. 4F). The cell death observed later on in the LPM is however not caused by the lack of *Fgf8* expression as it is also observed at non-limb levels. In contrast, paraxial mesoderm markers analysis shows that *Pax3* expression in the anterior PSM and early somites (Goulding et al., 1991) is normal (Fig. 6I,J). Similarly, *Fgf8* expression domains in the rostral and caudal lips of the dermomyotome (Crossley and Martin, 1995) are not affected at E9.5 in the mutant paraxial mesoderm (Fig. 6K,L). Expression of *Pax3* in the dermomyotome (Goulding et al., 1991) and of *Myf5* in the myotome (Ott et al., 1991) is however decreased in *T-Cre;Taf10* mutants (Fig. 6I,J,M,N). Delayed myotome formation was evidenced by immuno-localization of Myogenin or Myosin Heavy Chains at E9.5 and E10.5 (data not shown). Similar observations were made in *Hes7-Cre/+;Taf10*^flox/flox^ mutant embryos (Fig. S9). Expression of *Shh* in the notochord is normal (Echelard et al., 1993) indicating that the axial mesoderm is not obviously affected in the *T-Cre;Taf10* mutant embryos (Fig. 6O,P). Altogether, these results indicate different requirements for TAF10 depending on the type of mesoderm. However, we cannot rule out that the effect seen in the LPM arises secondarily to a defect in the developing paraxial mesoderm.

### Absence of TAF10 does not affect global steady state mRNA and cyclic transcription in the PSM

Our next goal was to investigate Pol-II transcription status in mutant embryos. We first compared steady state rRNA (Pol-I) and mRNA (Pol-II) transcript levels by quantifying the absolute expression levels of *RNA18S* versus classical Pol-II housekeeping genes (*Actc1*, *Gapdh* and *Rplp0*) (Fig. 7A). No significant differences between mutant and control samples were detected when comparing the results obtained with 3 different pairs of RNA 18S primers (Fig. 7B). The results are similar for *Gapdh* and *Rplp0* (Fig. 7B). Expression of the *Luvelu* reporter (Aulehla et al., 2008) in *T-Cre;Taf10* mutant embryos (Fig. 7C,D) supports the idea that cyclic transcription initiation still occurs in the *T-Cre;Taf10* mutant PSM at E9.5. Altogether these results indicate that, around E9.5, absence of detectable TAF10 does not affect global steady state mRNA and PSM-specific cyclic transcription.

**Fig. 7.**
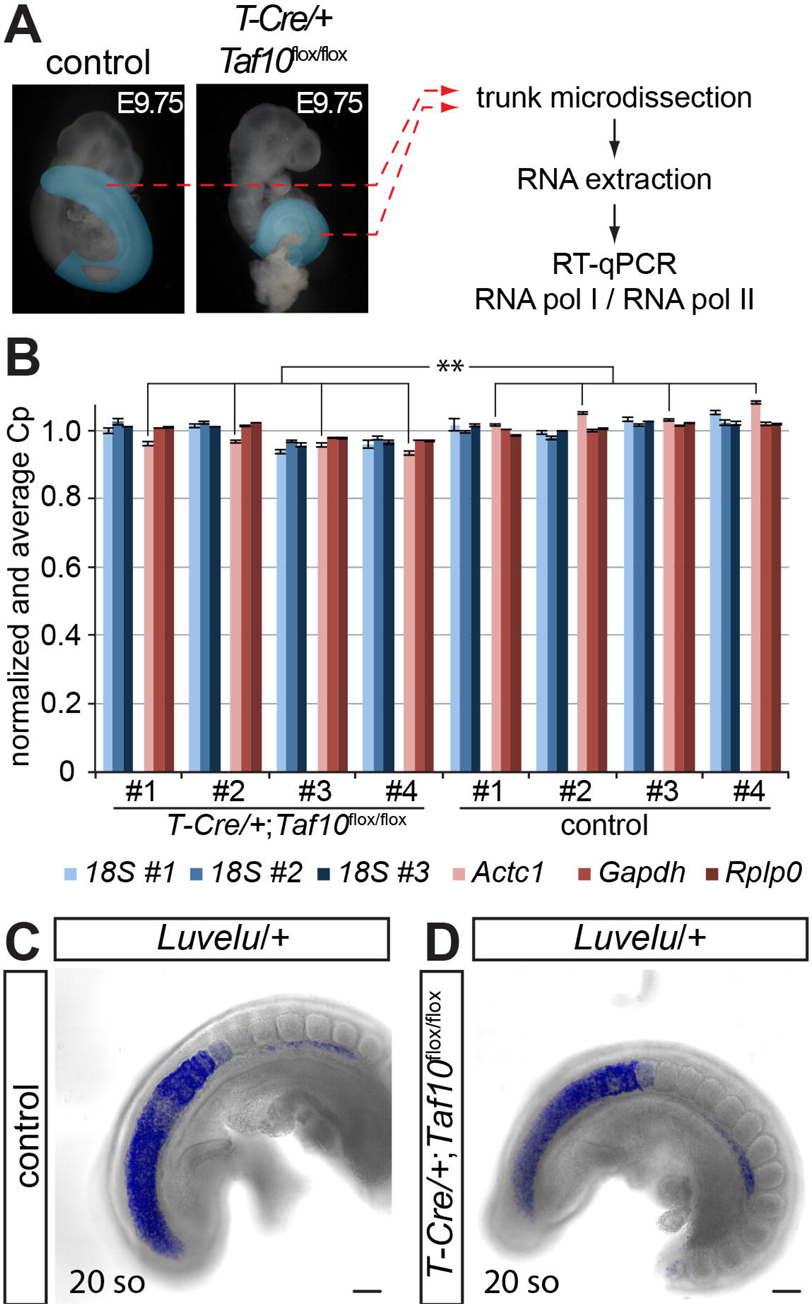
Global transcription is not affected in absence of TAF10 in the paraxial mesoderm. (A) Comparison between Pol-II and Pol-I transcription. Trunk axial structures highlighted in blue were dissected from E9.75 control and *T-Cre*/+;*Taf10*^flox/flox^ mutant embryos and RT-qPCR was performed for Pol-I and Pol-II specific housekeeping genes. (B) Comparison of averaged and normalized expression of Pol-I (blue) and Pol-II (red) specific markers from control (white) and mutant (grey) samples. **; p-value <0.01 (n=4, Aspin Welch corrected Student’s *t*-test). The error bars indicate s.e.m. (C,D) Confocal z-stack images projection of E9.5 *Luvelu/+* control © and *T-Cre*/+;*Taf10*^flox/flox^;*Luvelu/+* mutant (D). Scale bars in C-D represent 100 µm.

### Expression of specific genes is altered in the PSM at E9.5 in the absence of TAF10

We next performed a transcriptome analysis in order to see whether specific genes were affected in absence of TAF10. We performed microarray analyses from micro-dissected PSMs of E9.5 (17-19 somites) control and *T-Cre;Taf10* mutant embryos (Fig. 8A). Analysis by scatter plot shows that TAF10 loss has only very minor impact on gene expression (Fig. 8B). We then performed a statistical analysis using Fold Change Ranking Ordered Statistics (FCROS) (Dembélé and Kastner, 2014). We found 369 differentially expressed genes using a fold change cut-off of 1.5x (218 down-regulated and 151 up-regulated, see Table S2). This analysis identified genes related to the cell cycle, TAFs, signalling pathways, or *Hox*/*para-Hox* genes (see Table 1). We also observed that some genes previously identified as cyclic genes in the PSM such as *Egr1*, *Cyr61*, *Dkk1*, *Spry4* and *Rps3a* (Krol et al., 2011), are also differentially expressed in *T-Cre;Taf10* mutant PSMs (Table 1 and Fig. S10A). Interestingly, the highest up-regulated gene (4.8 fold) is *Cdkn1a* encoding a cyclin-dependent kinase inhibitor involved in G1 arrest (Dulić et al., 1994). We also identified *Gas5*, a tumor suppressor gene encoding 2 lncRNAs and several small nucleolar RNAs in its introns (Ma et al., 2015) as the highest down-regulated gene (from −2 to −4.9 fold). We confirmed the up-regulation of *Cdkn1a*, *Cdkn1c*, *Ccng1* and *Cdkl3*, and the down-regulation of *Gas5* by RT-qPCR using control and *T-Cre;Taf10* mutant tail tips (Fig. 8D). Up-regulation of *Cdkn1a* and *Cdkn1c* could explain the growth arrest that is observed in *T-Cre;Taf10* mutant embryos.

**Fig. 8.**
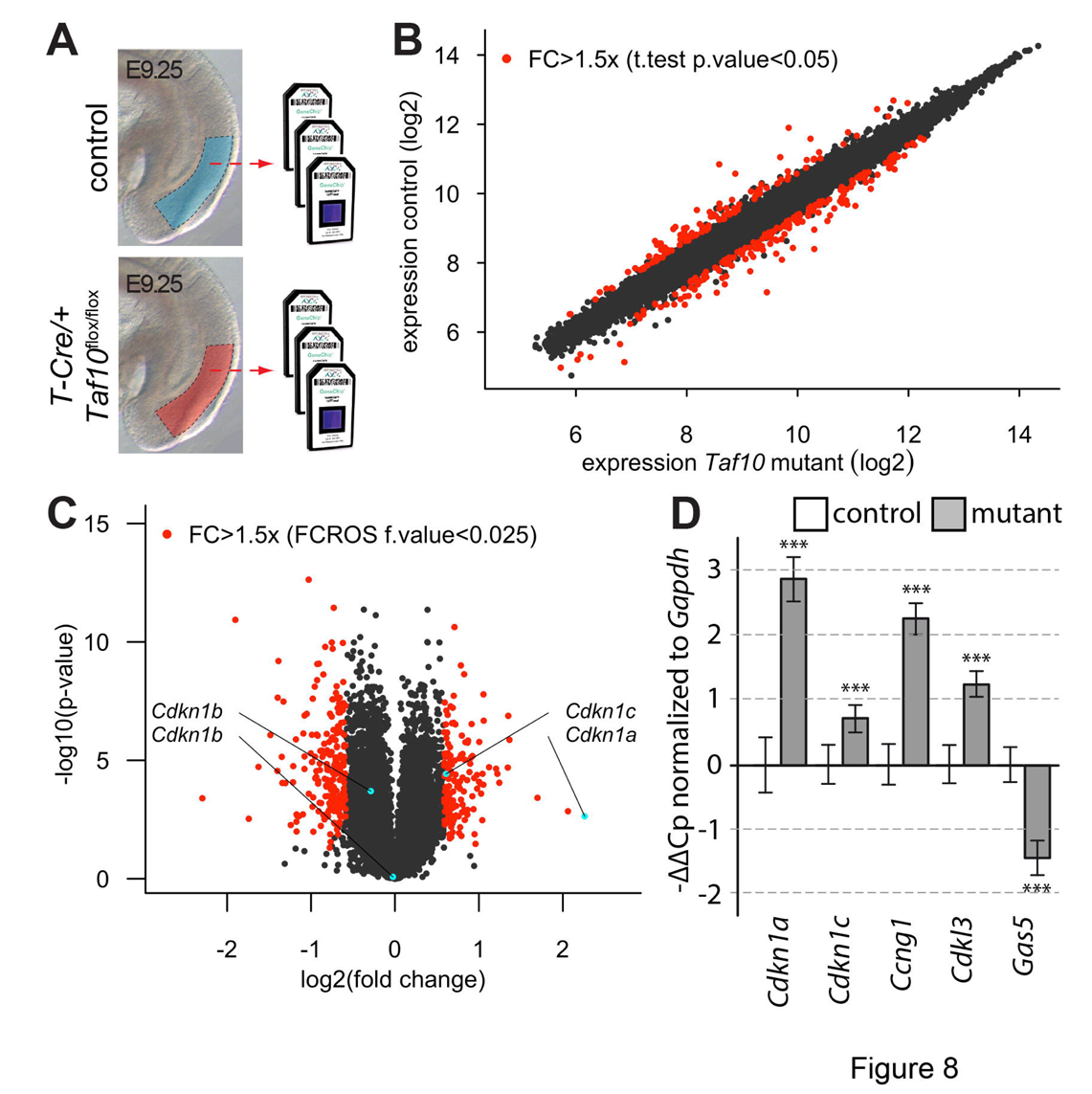
Limited specific effect on RNA polymerase II transcription in absence of TAF10 in the PSM. (A) Strategy used for the microarray analysis from E9.5 microdissected PSM of control (blue) and *T-Cre*;*Taf10* mutant (red) embryos. (B) Scatter plot comparing gene expression between control and *T-Cre*;*Taf10* mutant PSM. Red dots correspond to statistically significant differences for a fold change greater than 1.5 after a *t*-test. (C) Vulcano plot comparing gene expression between control and *T-Cre*/+;*Taf10*^flox/flox^ mutant PSM after FCROS analysis. Red dots correspond to statistically significant differences for a fold change greater than 1.5. (D) RT-qPCR analysis for cell-cycle genes candidates from E9.25 control (white) and *TCre*;*Taf10* mutant (grey) tail tips: -ΔΔCp are normalized to *Gapdh*. ns; non significant, *; p-value <0.05, **; p-value <0.01, ***; p-value <0.001 (n=4, Aspin Welch corrected Student’s *t*-test). The error bars indicate s.e.m.

**Table 1.** Selection of differentially expressed genes in the PSM of E9.5 *T-Cre;Taf10* mutant embryos. The statistical analysis has been performed using FCROS with a cut-off of 1.5 for the fold change. Difference is considered significant for a f value below 0.025 or above 0.975.

| description | symbol | absolute FC | f value |
| --- | --- | --- | --- |
| **cell cycle** |  |  |  |
| growth arrest specific 5 | *Gas5* | -4.9075408 | 0.017718801 |
| growth arrest specific 5 | *Gas5* | -3.736196532 | 0.017771121 |
| growth arrest specific 5 | *Gas5* | -2.634510323 | 0.017886397 |
| growth arrest specific 5 | *Gas5* | -2.072799706 | 0.018319525 |
| cyclin-dependent kinase inhibitor 1A (P21) | *Cdkn1a* | 4.789628182 | 0.982045384 |
| cyclin-dependent kinase inhibitor 1C (P57) | *Cdkn1c* | 1.525180853 | 0.979478088 |
| cyclin-dependent kinase-like 3 | *Cdkl3* | 1.779798823 | 0.981084679 |
| cyclin G1 | *Ccng1* | 2.005804019 | 0.981733355 |
| **RNA pol I associated complexes** |  |  |  |
| TATA box binding protein (Tbp)-associated factor, RNA polymerase I, D | *Taf1d* | -2.317074936 | 0.018087818 |
| TATA box binding protein (Tbp)-associated factor, RNA polymerase I, D | *Taf1d* | -2.266286542 | 0.01809225 |
| TATA box binding protein (Tbp)-associated factor, RNA polymerase I, D | *Taf1d* | -2.039738816 | 0.018555265 |
| TATA box binding protein (Tbp)-associated factor, RNA polymerase I, D | *Taf1d* | -1.785722753 | 0.019331127 |
| TATA box binding protein (Tbp)-associated factor, RNA polymerase I, D | *Taf1d* | -1.632420432 | 0.020404534 |
| **RNA pol II associated complexes** |  |  |  |
| TAF6 RNA polymerase II, TATA box binding protein (TBP)-associated factor | *Taf6* | 1.72440928 | 0.980946134 |
| TAF9B RNA polymerase II, TATA box binding protein (TBP)-associated factor | *Taf9b* | 1.591031326 | 0.978636615 |
| TAF5 RNA polymerase II, TATA box binding protein (TBP)-associated factor | *Taf5* | 1.536439612 | 0.978826464 |
| polymerase (RNA) II (DNA directed) polypeptide A | *Polr2a* | 1.5052891 | 0.979153598 |
| **signaling pathways and transcription factors** |  |  |  |
| Mix1 homeobox-like 1 (Xenopus laevis) | *Mixl1* | 1.565820524 | 0.97874697 |
| T-box 6 | *Tbx6* | 1.546625984 | 0.979948731 |
| E26 avian leukemia oncogene 2, 3' domain | *Ets2* | -1.537617023 | 0.021680906 |
| fibroblast growth factor 9 | *Fgf9* | -1.549855417 | 0.02153317 |
| ephrin A5 | *EfnA5* | -1.627750959 | 0.020793883 |
| dual specificity phosphatase 4 | *Dusp4* | -1.647697371 | 0.02022193 |
| R-spondin 3 homolog (Xenopus laevis) | *Rspo3* | -1.661619669 | 0.020011719 |
| cytochrome P450, family 26, subfamily a, polypeptide 1 | *Cyp26a1* | -1.671145052 | 0.020601729 |
| caudal type homeobox 4 | *Cdx4* | -1.51866511 | 0.023200463 |
| homeobox A7 | *Hoxa7* | 1.635575113 | 0.980606968 |
| homeobox B7 | *Hoxb7* | 1.822569059 | 0.981446929 |
| homeobox D1 | *Hoxd1* | 1.97112032 | 0.981663225 |
| homeobox A3 | *Hoxa3* | 2.549458032 | 0.981967438 |
| **cyclic genes** |  |  |  |
| early growth response 1 | *Egr1* | 1.610151831 | 0.979126918 |
| cysteine rich protein 61 | *Cyr61* | 1.713059121 | 0.981037055 |
| dickkopf homolog 1 (Xenopus laevis) | *Dkk1* | 1.944872688 | 0.981052429 |
| sprouty homolog 4 (Drosophila) | *Spry4* | -1.538775581 | 0.021864177 |
| ribosomal protein S3A | *Rps3a* | -1.585508682 | 0.020915584 |

Some TFIID-TAFs were also up-regulated: *Taf5* (1.5 fold), *Taf6* (1.7 fold) and *Taf9b* (1.6 fold) (Table 1, Table S2). We validated these differential expressions by RT-qPCR and found that the expression of most of the genes encoding the other TAFs were also up-regulated (Fig. S10B). The biological significance of these differences is not clear as no obvious increase in protein levels could be observed for TAF4A, TAF5 and TAF6 (Fig. 2A). *Taf10* expression is down-regulated in *T-Cre;Taf10* mutant tail tips and expression of *Taf8*, encoding TAF10’s main TFIID partner is also down-regulated in the mutants. These data suggest that the decreased level of TAF8 protein observed in *R26Cre;Taf10* lysates (Fig. 2A) could also be related to transcriptional regulation. No differences could be detected for the SAGA-specific TAF5L and TAF6L (Fig. S10C). Altogether, our data show that gene expression controlled by Pol-II is not initially globally affected in the absence of TAF10, however, the lack of TAF10 could induce a change in the steady state mRNA levels of specific genes.

## Discussion

The composition of TFIID and SAGA complexes in the developing mouse embryo has not yet been described. Here, we analysed the composition of these complexes in E9.5 mouse embryos in presence and absence of TAF10. We showed that the absence of TAF10 strongly affects TFIID and SAGA formation. *Taf10* deletion during somitogenesis confirmed the requirement of TAF10 during embryonic development in agreement with previous studies (Indra et al., 2005; Mohan et al., 2003; Tatarakis et al., 2008). However, in contrast to these studies, we identified a time window around E9.5 when no obvious somitogenesis defects are detected, despite the absence of detectable full-length TAF10 protein in mutant embryos. In these mutants, transcription is still broadly functional as shown by the lack of global effect on Pol-II transcription.

### TAF10 is required for TFIID and SAGA assembly during development

Our data demonstrate a global decrease in TFIID and of SAGA assembly in *Taf10* mutant embryos. In F9 cells, in absence of TAF10, TFIID is minimally affected by the release of TBP from the complex, while interaction between the different TAFs is maintained (Mohan et al., 2003) and in the liver, TFIID assembly is completely abrogated (Tatarakis et al., 2008). These differences could be explained either by cell-type specific differences or by difference in the timing of these analyses following *Taf10* deletion as Tatarakis *et al*. (2008) performed their experiments 8-15 days after *Taf10* deletion. The status of SAGA has not been previously investigated in *Taf10* mutant embryos. Our work demonstrates for the first time that not only TFIID, but also SAGA is affected in *Taf10* mutant embryos. Our new data show that the defect in the assembly of canonical TFIID and SAGA is observed already 2 days after the induction of *Taf10* deletion, a timing that coincides with the disappearance of detectable full length TAF10 protein. On the other hand, we can still detect reduced interactions between TAF7 and most of the TAFs following *Taf10* deletion suggesting that, as observed in HeLa or F9 cells, there could be some TFIID-like complexes that do not contain TAF10, albeit in reduced levels. Our data exclude the existence of similar TAF10-less SAGA-like complexes in the embryo.

TAF10 depletion is very efficient since no TAF10 proteins can be detected by western blot in the mutant embryo lysates. Analysis of the detected peptides strongly suggests that only in the TAF7-IP (TFIID), potential full-length TAF10 proteins are detected, albeit at very low frequency. This suggests that very low levels of canonical TFIID complexes could still be present at this stage. Furthermore, these results, in comparison with the SAGA-IPs, suggest that TAF10 is very stable when incorporated into TFIID, probably because of the lower rate of TFIID turnover compared to that of SAGA.

TFIID is built from sub-modules that assemble in the cytoplasm, at least *in vitro* (Bieniossek et al., 2013; Trowitzsch et al., 2015), and it is likely that such TFIID sub-modules are immunoprecipitated in our experiments since we performed our analyses using whole cell extracts. The TAF7 paralog; TAF7L, that has been associated with germ cells and adipocytes (Zhou et al., 2013a; Zhou et al., 2013b), is not present in TFIID-IPs, indicating that the majority of TFIID contains TAF7, at least at E9.5. However, other TAF paralogs, such as TAF4A and TAF4B, TAF9 and TAF9B are detected. This potential TFIID diversity could exist inside all the cells or could be cell specific and could explain the developmental differences observed between LPM and paraxial mesoderm. However, novel methods will be required to characterize the composition of TFIID and SAGA complexes in a cell type specific manner in the embryo.

### A TAF10 truncated protein can potentially be integrated into TFIID and SAGA submodules

Our strategy conditionally removes exon 2 and theoretically leads to the splicing of exon 1 to exon 3 (Mohan et al., 2003). These exons are not in frame and therefore, the 77 amino-acids coded by exon 1 are followed by 15 extra amino-acids in the mutant (Fig. S4D). This mutant protein has the N-terminal unstructured domain of TAF10 but more importantly, lacks its HFD required for the interaction with TAF3, TAF8 or SUPT7L/ST65G (Soutoglou et al., 2005). HFD-HFD interactions are crucial for nuclear import of TFA10 which does not contain any NLS (Soutoglou et al., 2005). Since no TFIID subunits could be co-immunoprecipitated from whole cell extracts of *R26*^CreERT2/R^;*Taf10*^flox/flox^ ES cells after 4-hydroxy tamoxifen treatment with an antibody that recognizes the N terminal part of TAF10 (Fig. S5), it is very unlikely that this truncated protein can be incorporated in mature SAGA and TFIID complexes that are functional in the nucleus. However, we cannot rule out that this truncated protein could incorporate into rare cytoplasmic sub-modules containing TAF7 or SUPT3. Nevertheless, because the *Taf10* mutant heterozygotes are indistinguishable from control embryos (Fig. 1A), this also argues against a dominant negative effect of this peptide.

Another interesting question is the functionality of these potentially partial TFIID and/or SAGA complexes fully depleted of TAF10 protein or containing the truncated TAF10 protein. From our data, it is obvious that these different partial complexes cannot fully compensate for the loss of wild type complexes, but one cannot rule out a partial activity. Future analyses of the difference between the different types of mesoderm could help to elucidate whether such partial non-canonical TFIID and/or SAGA complexes have activities.

### Differential sensitivity to Taf10 loss in the mesoderm

*Taf10* deletion in the mesoderm or in the whole embryo leads to growth arrest that could be explained by the up-regulation of *Cdkn1a* and *Cdkn1c* expression. Similar observations were made in yeast (Kirschner et al., 2002) and in F9 cells (Metzger et al., 1999), following depletion of TAF10. Surprisingly, we also observed the down regulation of the tumour suppressor *Gas5* that is associated with increased proliferative and anti-apoptosis effects in cancer cells (Pickard and Williams, 2015). Interestingly, *Cdkn1a* expression is positively controlled by *Gas5* in stomach cancer at the transcript and protein levels (Liu et al., 2015). It is thus possible that TAF10 is required for the correct functioning of the *Gas5* regulatory network during development.

The phenotypes of null mutations of TFIID-TAFs coding genes such as *Taf7* (Gegonne et al., 2012) or *Taf8* (Voss et al., 2000) are very similar to that of the *Taf10* mutant phenotype (Mohan et al., 2003). In particular, these mutations are embryonic lethal around implantation stage. Moreover, *Taf7* null MEFs stop proliferating, suggesting that the growth arrest observed in our mutants is a direct consequence of the failure to properly build TFIID. We cannot exclude a potential contribution of SAGA loss in our mutants. However, deletion of genes coding for different enzymatic activities of SAGA such as *Kat2a*;*Kat2b* or *Usp22* are embryonic lethal, but with phenotypes much less severe than *Taf10* mutation (Lin et al., 2012; Xu et al., 2000; Yamauchi et al., 2000). Interestingly, axial and paraxial mesoderm formation is affected in *Kat2a*;*Kat2b* mutants whereas, extra-embryonic and cardiac mesoderm formation is not (Xu et al., 2000), strongly suggesting that SAGA could also have different functions in different types of mesoderm.

Another striking observation is that, while no TAF10 protein could be detected as early as E8.5 in the mesoderm of *T-Cre;Taf10* mutant embryos, we observed a difference of sensitivity to *Taf10* loss between the LPM (and the intermediate mesoderm) and the paraxial mesoderm. Interestingly, we observed a very early defect in the LPM with strong down-regulation of specific markers and absence of limb bud outgrowth. The absence of limb buds could be explained by a defect in FGF10 signalling activation in the mesoderm and/or by cell death in the LPM that occurs earlier than in the paraxial mesoderm of *T-Cre;Taf10* mutants. The relative resistance of the mutant paraxial mesoderm to cell death also suggests a difference of sensitivity. A similar observation has been made in F9 cells where RA induced differentiation of F9 cells into primitive endoderm rescued the apoptosis of *Taf10* mutant cells (Metzger et al., 1999). Interestingly, this effect was not observed when F9 cells are differentiated into parietal endoderm in the presence of RA and cAMP (Metzger et al., 1999). One interesting possibility could be that, being the principal source of RA (Niederreither et al., 1997), the paraxial mesoderm is protected from cell death in the mutant embryos via an autocrine mechanism. Another difference of sensitivity has also been observed in *Taf10* mutant blastocysts where inner cell mass dies by apoptosis whereas trophoblast can be maintained in culture (Mohan et al., 2003). It is interesting to note that trophoblast, primitive and parietal endoderms are extra-embryonic structures and are not part of the fully-developed embryo. The difference of sensitivity to the loss of *Taf10* in the mesoderm is the first to be observed *in vivo*, in an embryonic lineage. Since *Taf10* was deleted in paraxial mesoderm and LPM progenitors, we cannot rule out that the LPM increased sensitivity is indirect and mediated by the paraxial mesoderm, although we did not observe any obvious change in gene expression in the PSM at a time when limb bud development is already affected. Nevertheless, a tempting speculation is that TAF10 could serve as an interface of interaction with a LPM specific transcription factor as it has been described recently for GATA1 during erythropoiesis (Papadopoulos et al., 2015).

## Material and Methods

### Mice

Animal experimentation was carried out according to animal welfare regulations and guidelines of the French Ministry of Agriculture (ethical committee C2EA-17 projects 2012-077, 2012-078, 2015050509092048). All the lines have already been described (see supplementary methods). The day of vaginal plug was scored as embryonic day (E)0.5. Tamoxifen (Sigma) resuspended at 20 mg/ml in 5% EtOH/filtered sunflower seed oil was injected intraperitoneally (150 µl (3 mg) for a 20 g mouse) at E7.5.

### Embryos whole cell extracts

E9.5 mouse embryos (16-20 somites) were lysed in 10% glycerol, 20 mM Hepes (pH7), 0.35 M NaCl, 1.5 mM MgCl2, 0.2 mM EDTA, 0.1% Triton X-100 with protease inhibitor cocktail (PIC, Roche) on ice. Lysates were treated 3 times with pestle stroke followed by 3 liquid nitrogen freezing-thaw cycles. Lysates were centrifuged at 20817 rcf for 15 min at 4°C and the supernatants were used directly for IPs or stored at −80°C for western blots.

### Immunoprecipitations

Inputs were incubated with Dynabeads coated with antibodies (see supplementary methods and Table S3) overnight at 4°C. Immunoprecipitated proteins were washed twice 5 min with 500mM KCl buffer (25 mM Tris-HCl (pH7), 5 mM MgCl2, 10% glycerol, 0.1% NP40, 2 mM DTT, 500 mM KCl and PIC (Roche)), then washed twice 5 min with 100 mM KCl buffer (25 mM Tris-HCl (pH7), 5 mM MgCl2, 10% glycerol, 0.1% NP40, 2 mM DTT, 100 mM KCl and PIC (Roche)) and eluted with 0.1 M glycine (pH2.8) for 5 min three times. Elution fractions were neutralized with 1.5 M Tris-HCl (pH8.8).

### Western blots

Immune complexes or 15 µg of embryo lysates were boiled 10 min in 100 mM Tris-pH6.8, 30% glycerol, 4% SDS, 0.2% bromophenol blue, 100 mM DTT, resolved on precast SDS-polyacrylamide gel 4-12% (Novex) and transferred to nitrocellulose membrane (Protran, Amersham). Membranes were blocked in 3% milk in PBS for 30 min and incubated with the primary antibody (Table S3) overnight at 4°C. Membranes were washed three times 5 min with 0.05% Tween20 in PBS. Membranes were incubated with HRP-coupled secondary antibodies for 50 min at RT, followed by ECL detection (ThermoFisher).

### Mass spectrometry analyses

Samples were analyzed using an Ultimate 3000 nano-RSLC (Thermo Scientific, San Jose, California) coupled in line with a linear trap Quadrupole (LTQ)-Orbitrap ELITE mass spectrometer via a nano-electrospray ionization source (Thermo Scientific). Data were analyzed by calculation of the NSAFbait (see supplementary methods).

### Section and Immunolocalization

Embryos were fixed in 4% PFA 2 hours at 4°C, rinsed 3 times in PBS, equilibrated in 30% sucrose/PBS and embedded in Cryomatrix (Thermo Scientific) in liquid nitrogen vapours. Twenty µm sections were obtained on a Leica crysotat. Immunolabelling was performed as described (Vincent et al., 2014). Sections were counterstained with DAPI (4′,6-diamidino-2-phenylindole, dihydrochloride, Molecular Probes) and imaged with a LSM 510 laser-scanning microscope (Carl Zeiss MicroImaging) through a 20x Plan APO objective (NA 0.8).

### Luvelu imaging

Freshly dissected embryos were kept in DMEM without red phenol (Life technologies). Luvelu signal was detected using a SP5 TCS confocal microscope (Leica) through a 20x Plan APO objective (NA 0.7).

### Whole-mount in situ hybridization (WISH), X-gal and Lysotracker Red staining

WISH were performed as described (Nagy et al.). *Axin2*, *Fgf8*, *Hand2*, *Lfng*, *Msgn1*, *Myf5*, *Shh*, *Snai1*and *Uncx4.1* probes have been described (Aulehla and Johnson, 1999; Aulehla et al., 2008; Crossley and Martin, 1995; Dale et al., 2006; Echelard et al., 1993; Mansouri et al., 1997; Ott et al., 1991; Srivastava et al., 1997; Yoon et al., 2000). A minimum of 3 embryos were used for the classical markers and a minimum of 7 were used for the cyclic genes. X-gal and Lysotracker Red (Molecular Probes) stainings were performed as described (Rocancourt et al., 1990; Vincent et al., 2014).

### RT-qPCR and statistical analysis

Micro-dissected embryo tail tip or trunk tissue (without limb buds for the controls) were lysed in 500 µl TRIzol (Life technologies). RNA was extracted according to the manufacturer’s recommendations and resuspended in 20 µl (trunk) or 11 µl (tail tips) of RNase free water (Ambion). Reverse transcription was performed using the QuantiTect Reverse Transcription Kit (Qiagen) in 12 µl reaction volume and diluted by adding 75 µl of RNase free water. Quantitative PCRs were performed on a Roche LightCycler II 480 using LightCycler 480 SYBR Green I Master (Roche) in 8 µl reaction (0.4 µl cDNA, 0.5 µM primers). Four mutants and four controls with the same somite number were analysed in triplicates. Statistical analysis and primer sequences are described in the supplementary methods and Table S4.

### Microarrays and statistical analysis

Posterior PSM of E9.5 embryos were individually microdissected (Dequéant et al., 2006) and lysed in 200µl TRIzol (Life technologies), and yolk sac was used for genotyping. Three PSM of 17-19 somites embryos of the same genotype were pooled for one replicate and analysed on GeneChip^®^ MoGene1.0ST (Affymetrix). Data were normalized using RMA (Bioconductor), filtered and FCROS (Dembélé and Kastner, 2014) was used for the statistical analysis (see supplementary methods).

## Acknowledgements

We thank Violaine Alumni and the Biochip and Sequencing platform (IGBMC) for the microarray experiments, Doulaye Dembele for his advices on the statistical analysis of the microarrays, Mathilde Decourcelle and the Proteomic platform (IGBMC) for the Orbitrap analyses. We also thank Eli Scheer for her skilful advices on immunoprecipitations, Ivanka Kamenova for her help to validate the antibodies and Joël Herrmann for the validation of the qPCR primers. We thank Didier Devys, Goncalo Vilhais-Neto and Ziad Al Tanoury for their critical reading of the manuscript.

## Author contributions

PB performed, interpreted the experiments and contributed to the writing of the manuscript. AH contributed to the imaging and to the *Luvelu* experiments and MF initiated the mass spectrometry analyses. MJ performed the MS analyses. OP and LT funded, contributed to the planning and interpretation of experiments, and contributed to the writing of the manuscript. SDV supervised, planned, performed, interpreted the experiments and wrote the manuscript.

## Data availability

Raw microarray data were deposited in GEO database (GSE82186). Raw mass spectrometry data are available via ProteomeXchange (PXD004688).

